# *In vitro* and *in vivo* roles of glucocorticoid and vitamin D receptors in the control of cardiomyocyte proliferative potential

**DOI:** 10.1101/755827

**Authors:** Stephen Cutie, Dominic Lunn, Guo N. Huang

## Abstract

Cardiomyocyte (CM) proliferative potential varies considerably across species. While lower vertebrates and neonatal mammals retain robust capacities for CM proliferation, adult mammalian CMs lose proliferative potential due to cell-cycle withdrawal and polyploidization, failing to mount a proliferative response to regenerate lost CMs after cardiac injury. The decline of murine CM proliferative potential occurs in the neonatal period when the endocrine system undergoes drastic changes for adaptation to extrauterine life. We recently demonstrated that thyroid hormone (TH) signaling functions as a primary factor driving CM proliferative potential loss in vertebrates. Whether other hormonal pathways govern this process remains largely unexplored. Here we showed that agonists of glucocorticoid receptor (GR) and vitamin D receptor (VDR) suppressed neonatal CM proliferation *in vitro*. We next examined CM nucleation and proliferation in mutant mice lacking GR or VDR specifically in CMs, but we observed no difference between mutant and control littermates. Additionally, we generated compound mutant mice that lack GR or VDR and express dominant-negative TH receptor alpha in their CMs, and similarly observed no increase in CM proliferative potential compared to dominant-negative TH receptor alpha mice alone. Thus, although GR and VDR activation in cultured CMs is sufficient to inhibit CM proliferation, they seem to be dispensable for CM cell-cycle exit and binucleation *in vivo*. In addition, given the recent report that VDR activation in zebrafish promotes CM proliferation and tissue regeneration, our results suggest distinct roles of VDR in zebrafish and rodent CM cell-cycle regulation.

## BACKGROUND

Cardiovascular disease is the leading killer in the United States, mostly due to heart failure after myocardial infarction (MI)^1,2^. After ischemic injury like MI induces CM death in the hearts of adult mammals, lost CMs are not replenished and fibrotic tissue permanently replaces previously functional cardiac muscle. However, lower vertebrates like zebrafish (*Danio rerio*), newts (*Notophthalmus viridescens*), and axolotls (*Ambystoma mexicanum*) display robust CM proliferation and myocardial regeneration after cardiac injury^2–4^. Intriguingly, neonatal mammals also transiently possess considerable CM proliferative potential. Ischemic heart injury induced in newborn mice at P0 or P1 resolves as fibrosis-free, fully-regenerated cardiac muscle by 21 days post-injury, mediated by existing CMs that proliferate to reconstitute the lost myocardium^5,6^. This transient CM proliferative potential is lost in mice after the first week as neonatal CMs binucleate and permanently exit the cell cycle^7^. Because the regeneration-competent hearts of lower vertebrates and neonatal mammals consist predominantly of mononuclear diploid CMs while the non-regenerative hearts of adult mammals consist primarily of polyploid CMs, this developmental polyploidization – documented in both mice and humans – is implicated as a critical inhibitor of CM proliferative potential^4,8–10^. However, the underlying molecular processes that drive CM polyploidization and cell cycle withdrawal are not fully understood.

Recent evidence indicates that thyroid hormones (THs) play a crucial role in driving CM polyploidization and suppressing CM proliferative potential in vertebrates^11^. TH signaling is a critical regulator of metabolism and thermogenesis conserved across vertebrates and is particularly active in endotherms^12^. THs enhance CM contractility *in vivo* and rise soon after birth in neonatal mice, coinciding with the closure of the CM proliferative window^13,14^. Serum TH levels are also substantially lower in newts and zebrafish than in non-regenerative mammals^15,16^. Neonatal mice dosed with propylthiouracil (PTU) – a potent inhibitor of TH synthesis – from birth through postnatal day 14 (P14) show significantly increased mononuclear CM percentage and CM proliferation^11^. These mononuclear CMs and CM proliferation phenotypes are also observed in mutant mice in which TH signaling is specifically inactivated in CMs^11^. These mutant mouse hearts also display enhanced recovery after MI as indicated by cardiac contractile functions and fibrosis one month post-surgery. Conversely, exogenous TH suppresses CM proliferation and cardiac regeneration in adult zebrafish. These data strongly suggest an inhibitory role played by TH in the developmental control of CM proliferative potential.

In addition to TH, other hormones such as glucocorticoids and vitamin D have been shown to suppress CM proliferation in culture. Glucocorticoids are stress-associated steroid hormones produced by the adrenal glands that act on nearly all organs in the body via binding to the glucocorticoid receptor (GR)^17,18^. Specifically, glucocorticoids like dexamethasone are established cell cycle regulators known to repress the cell cycle primarily through the GR, which acts as a transcription factor after glucocorticoid binding^19,20^. It has been reported that GR activation inhibits neonatal rat CM proliferation and increases CM binucleation through epigenetic repression of Cyclin D2 gene^21,22^. Furthermore, adult zebrafish heart regeneration is impaired by either GR agonist exposure^23^ or crowding-induced stress through GR activation^24^.

Vitamin D is a steroid hormone precursor that is hydroxylated sequentially in the liver and kidneys into ercalcitriol (D2) or calcitriol (D3). These active hormones can bind to vitamin D receptors (VDRs) and regulate downstream gene expression in target cells^25^. Alfacalcidol (Alfa) is a vitamin D analog that forms calcitriol directly after hydroxylation in the liver^26^. Anti-proliferative effects of vitamin D analogs have been reported in mammalian cells generally^27^ and mammalian CMs and CM-derived cell lines specifically^28,29^. However, vitamin D analogs alfacalcidol and calcipotriene significantly increase CM proliferation in embryonic zebrafish, whereas VDR-inhibitor PS121912 significantly decreases CM proliferation^30^. Alfacalcidol also stimulates CM proliferation in adult zebrafish during heart regeneration, while VDR suppression decreases CM proliferation during regeneration^30^.

Intriguingly, CM-specific deletion of either GR or VDR results in cardiac hypertrophy^31,32^. This cardiac enlargement phenotype together with the documented functions to inhibit CM proliferation *in vitro* led us to investigate the possibility that GR and VDR signaling alters CM proliferative potential *in vivo* either alone or in combination with TH signaling activation.

## RESULTS

We first examined if GR and VDR activation regulates CM proliferation *in vitro*. Primary CMs were isolated from neonatal mice and cultured with three GR agonists hydrocortisone, corticosterone, and dexamethasone for 48 hours. Proliferating CM nuclei were positive for DAPI (blue), PCM-1 rings (green), and EdU incorporation (red) (Fig. 1). CM proliferation is inhibited by hydrocortisone (0.30 ± 0.03%), corticosterone (0.15 ± 0.1%), and dexamethasone (0.07 ± 0.09%) as compared to CMs cultured without any GR agonists (1.22 ± 0.44%) (Fig. 1A). We also cultured primary neonatal mouse CMs with two VDR agonists: calcitriol and alfacalcidol. Neonatal CM proliferation was inhibited by 1 μM calcitriol (0.35 ± 0.07%) and 1 μM alfacalcidol (0.40 ± 0.02%) relative to CMs cultured without any VDR agonists (1.11 ± 0.11%) (Fig. 1B). This *in vitro* suppression of CM proliferation led us to investigate the *in vivo* contribution of GR and VDR to cardiomyocyte proliferative potential.

**Fig. 1.**
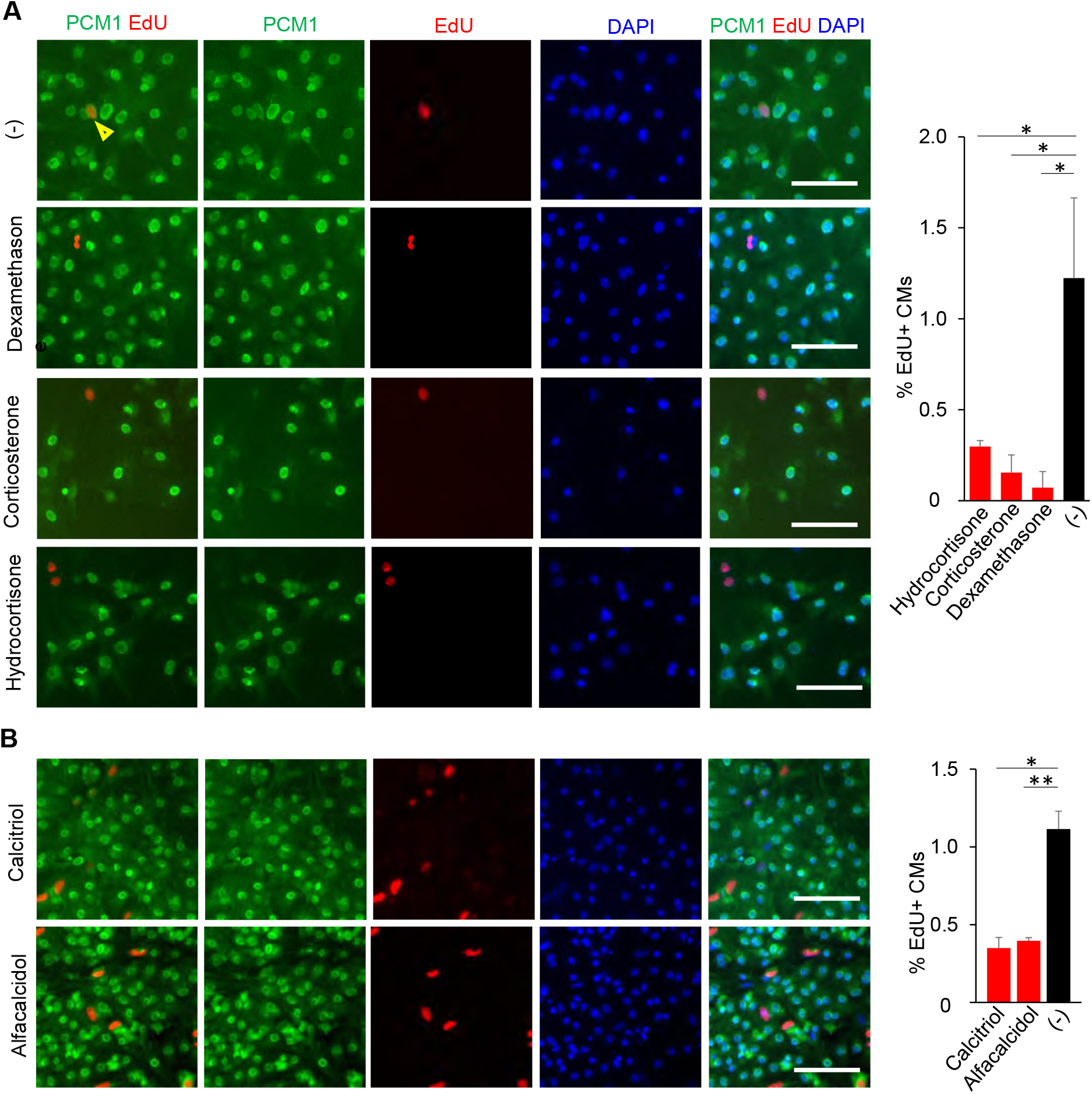
Glucocorticoid receptor (GR) and vitamin D receptor (VDR) agonists suppress cardiomyocyte (CM) proliferative potential *in vitro*. (**A**) Primary CMs cultured from P1 neonatal mice in the presence of 100 nM hydrocortisone, 100 nM corticosterone, or 100 nM dexamethasone incorporated 5- to 10-fold less EdU over 48 hr than CMs cultured without any GR agonist (−). (**B**) Similarly, P1 primary CMs cultured with 1 μM calcitriol and 1 μM alfacalcidol incorporated 2.5-fold less than those cultured with no VDR agonist. Values are reported as mean ± SEM (n=3). NS, not significant. \**p* < 0.05, \*\**p* < 0.01. Scale: 100 μm.

To elucidate the physiological role of GR in regulating CM proliferative potential *in vivo*, we generated *Myh6-Cre;Gr*^*f/f*^ knockout mice in which CM-specific CRE recombinase (Myh6-Cre) deletes the *Gr* gene (also named as Nuclear Receptor Subfamily 3 Group C member 1, *Nr3c1*), resulting in CM-specific loss of GR signaling (Fig. 2A). We examined heart phenotypes at P14 when mouse CMs have completed binucleation and withdrawn from the cell cycle^7^. Interestingly, unlike our *in vitro* primary CMs, no changes of CM proliferative activity *in vivo* were observed at P14 (Fig. 2B-D). *Myh6-Cre*;*Gr*^*f/f*^ knockout mice did not show an increase in the heart weight (mg) to body weight (g) ratio (6.21 ± 0.16) compared to littermate controls (6.16 ± 0.18) (Fig. 2B). Additionally, P14 *Myh6-Cre*;*Gr*^*f/f*^ hearts were comparable to littermate control hearts in terms of mononuclear CM percentage (13.6 ± 1.1% vs 13.7 ± 2.0%) (Fig. 2C). Analysis of CM proliferation using the Ki67 marker also showed a no increase in Ki67^+^ CMs in mutant hearts (2.1 ± 0.7% vs 2.2 ± 0.3%) (Fig. 2D). Altogether, we did not observe that CM-specific loss of GR affects CM proliferative potential.

**Fig. 2.**
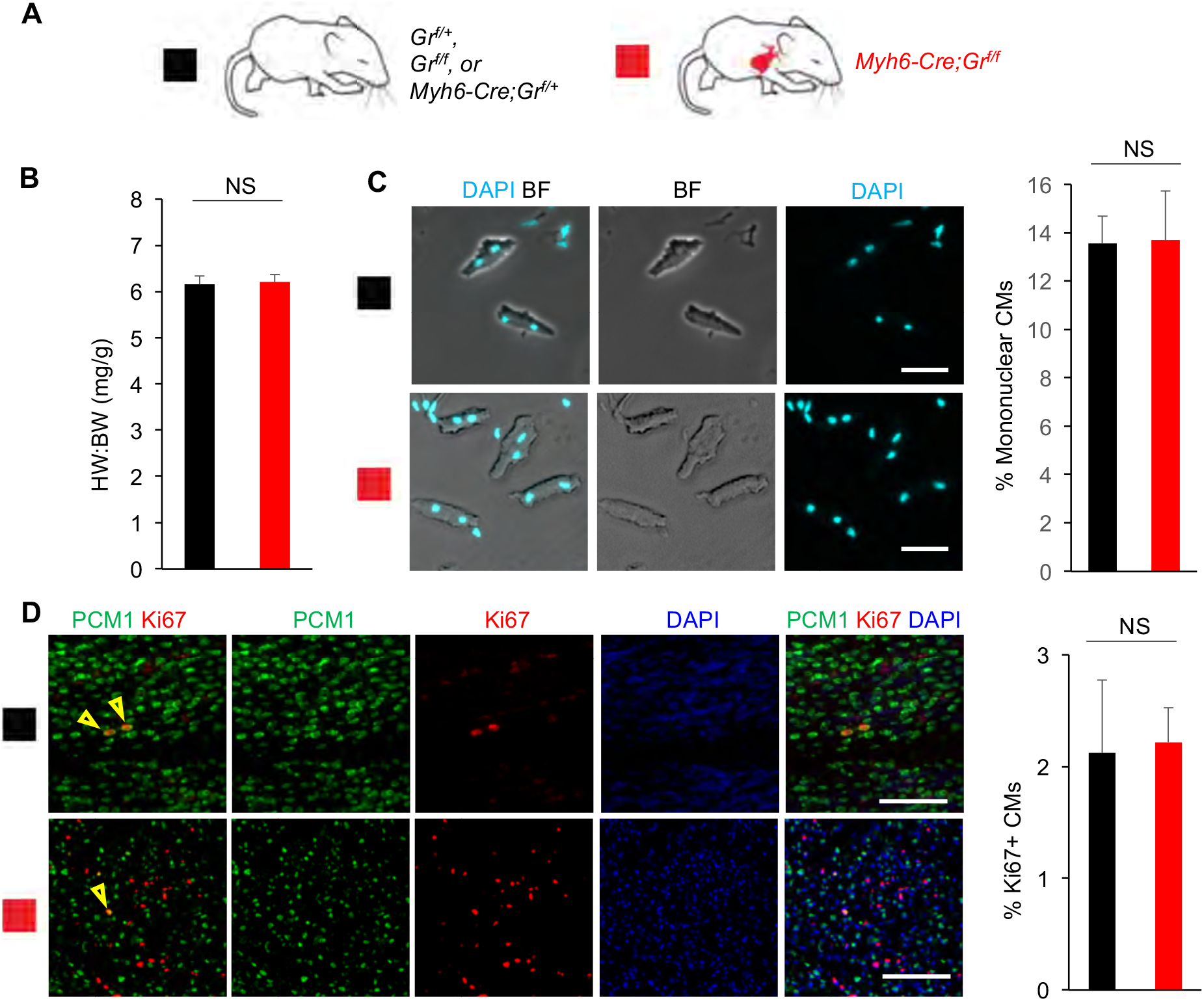
CM-specific loss of Gr signaling does not enhance CM proliferative potential *in vivo*. (**A**) Schematic for generating *Myh6-Cre;Gr*^*f/f*^ mice with CM-specific deletion of floxed endogenous GR and assessing CM proliferative potential at P14. Littermate controls included the following phenotypically-identical genotypes: *Gr*^*f/f*^, *Gr*^*f*/+^, and *Myh6-Cre;Gr*^*f*/+^. (**B**) Heart weight (mg) to body weight (g) ratio (HW:BW) did not differ between *Myh6-Cre;Gr*^*f/f*^ mice and littermate controls at P14. (**C**) Mononuclear CM percentage did not differ between *Myh6-Cre;Gr*^*f/f*^ mice and littermate controls at P14. (**D**) No difference in the number of Ki67^+^ CMs was observed between *Myh6-Cre;Gr*^*f/f*^ mice and littermate controls at P14. Values are reported as mean ± SEM (n=3). NS, not significant. \**p* < 0.05. Scale: (C) 50 μm (D) 100 μm.

To investigate the contribution of VDR to the control CM proliferative potential *in vivo*, we generated mice with CM-specific loss of VDR (*Myh6-Cre*;*Vdr*^*f/f*^) (Fig. 3A). As with GR deletion, no changes in CM proliferative activity were observed at P14 (Fig. 3B-D). The heart weight to body weight ratio did not significantly differ between *Myh6-Cre*;*Vdr*^*f/f*^ knockout mice (7.17 ± 0.76) compared to littermate controls (6.83 ± 0.57) (Fig. 3B). Additionally, P14 *Myh6-Cre*;*Vdr*^*f/f*^ hearts did not significantly differ from littermate control hearts in terms of mononuclear CM percentage (15.8 ± 3.2% vs 14.0 ± 1.0%) or CM proliferation (0.9 ± 0.3% vs 0.6 ± 0.1%) (Fig. 3C,D).

**Fig. 3.**
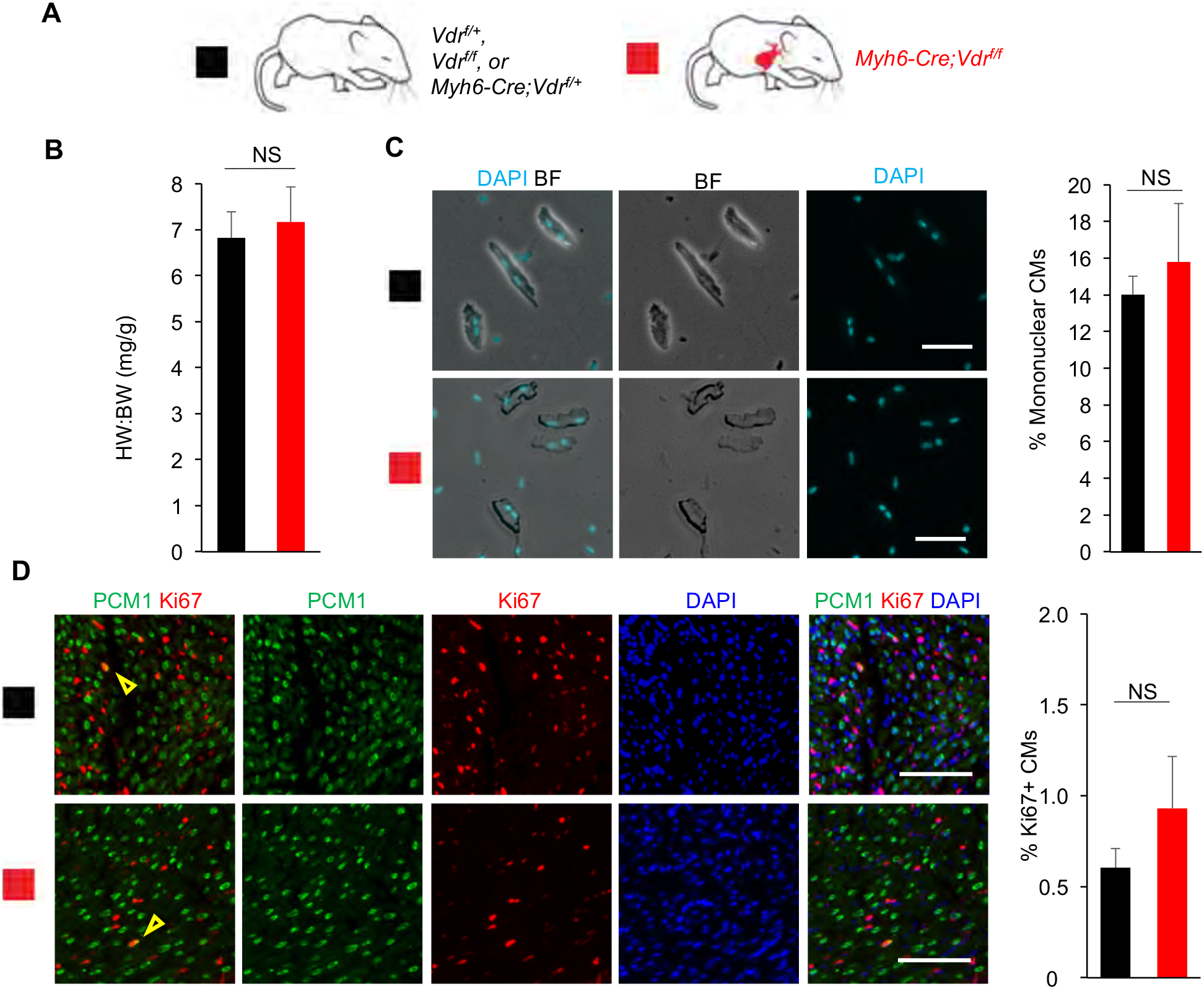
CM-specific loss of Vdr signaling does not enhance CM proliferative potential *in vivo*. (**A**) Schematic for generating *Myh6-Cre;Vdr*^*f/f*^ mice with CM-specific deletion of floxed endogenous VDR and assessing CM proliferative potential at P14. Littermate controls included three phenotypically-identical genotypes: *Vdr*^*f/f*^, *Vdr*^*f*/+^, and *Myh6-Cre;Vdr*^*f*/+^. (**B**) Heart weight (mg) to body weight (g) ratio (HW:BW) did not differ between *Myh6-Cre;Vdr*^*f/f*^ mice and littermate controls at P14. (**C**) Mononuclear CM percentage did not differ between *Myh6-Cre;Vdr*^*f/f*^ mice and littermate controls at P14. (**D**) No difference in the number of Ki67^+^ CMs was observed between *Myh6-Cre;Vdr*^*f/f*^ mice and littermate controls at P14. Values are reported as mean ± SEM (n=3). NS, not significant. \**P* < 0.05. Scale: (C) 50 μm (D) 100 μm.

We recently showed that TH signaling regulates CM proliferative potential, but mice with deficiency in TH receptor activation in CMs have ~30% mononuclear CMs at P14, implicating the existence of other pathways that contribute to CM binucleation in the neonatal period^11^. We next investigated whether loss of GR or VDR signaling affect CM proliferative potential in combination with loss of TH signaling. We bred *Myh6-Cre*;*Thra*^DN/+^;*Gr*^*f/f*^ mice that express dominant negative TH receptor alpha (THRA^DN^) and delete *Gr* specifically in CMs (Fig. 4A). The heart weight to body weight ratio for *Myh6-Cre*;*Thra*^DN/+^;*Gr*^*f/f*^ (7.28 ± 0.05) did not significantly differ from littermate *Myh6-Cre;Thra*^DN/+^;*Gr*^*f*/+^ controls (7.61 ± 0.55) (Fig. 4B). Additionally, compound mutant hearts did not display significant increases in CM proliferation (3.5 ± 0.7% vs 3.2 ± 0.7%) or CM nucleation (32.7 ± 8.7% vs 33.4 ± 6.1%) compared to littermate *Myh6-Cre;Thra*^DN/+^;*Gr*^*f*/+^ controls at P14 (Fig. 4C,D). Thus, we did not observe any significant effect of GR loss on top of TH signaling suppression.

**Fig. 4.**
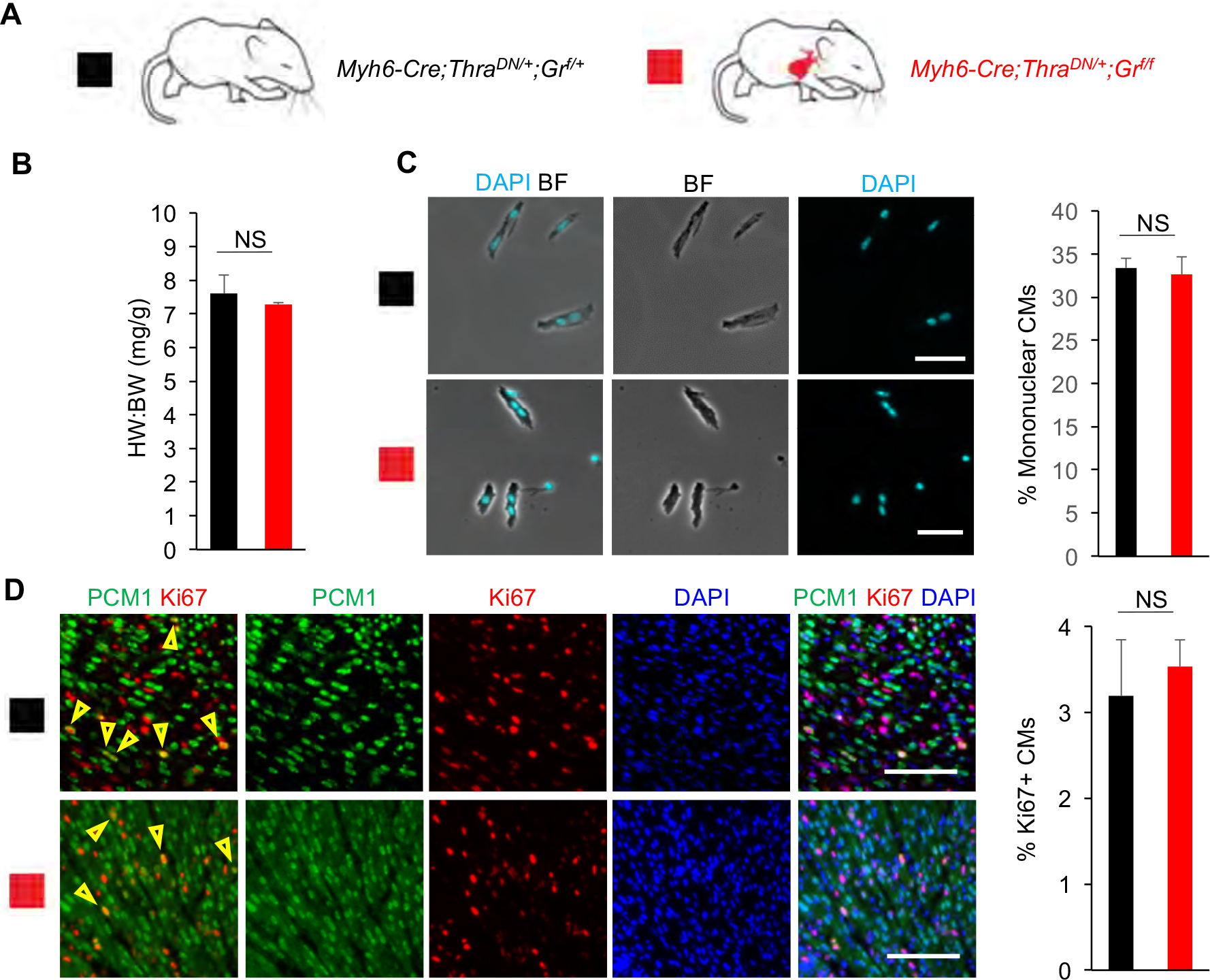
Loss of Gr is insufficient to alter the *in vivo* proliferative potential retained by thyroid hormone (TH) signaling-deficient CMs. (**A**) Schematic for generating *Myh6-Cre;Thra*^*DN*/+^;*Gr*^*f/f*^ mice with CM-specific deletion of floxed endogenous GR and CM-restricted expression of a dominant negative (DN) TH receptor alpha. (**B**) Heart weight (mg) to body weight (g) ratio (HW:BW) did not differ between *Myh6-Cre;Thra*^*DN*/+^;*Gr*^*f/f*^ mice and *Myh6-Cre;Thra*^*DN*/+^;*Gr*^*f*/+^ controls at P14. (**C**) Mononuclear CM percentage did not differ between *Myh6-Cre;Thra*^*DN*/+^;*Gr*^*f/f*^ mice and *Myh6-Cre;Thra*^*DN*/+^;*Gr*^*f/f*^ controls at P14. (**D**) No difference in the number of Ki67^+^ CMs was observed between *Myh6-Cre;Thra*^*DN*/+^;*Gr*^*f/f*^ mice and *Myh6-Cre;Thra*^*DN*/+^;*Gr*^*f/f*^ controls at P14. Values are reported as mean ± SEM (n=3). NS, not significant. \**p* < 0.05. Scale: (C) 50 μm (D) 100 μm.

Finally, we bred *Myh6-Cre;Thra*^DN/+^;*Vdr*^*f/f*^ mice and observed no significant difference in the heart weight to body weight ratio (6.83 ± 0.14 vs 7.23 ± 0.83), CM nucleation (24.4 ± 2.3% vs 26.9 ± 3.4%), or CM proliferation (3.0 ± 0.3 vs 3.9 ± 1.0%) compared to *Myh6-Cre;Thra*^DN/+^;*Vdr*^*f*/+^ littermate controls (Fig. 5A-D). As such, we did not observe a significant effect of VDR loss on CM proliferative potential on top of TH signaling suppression. Overall, our results show that although GR and VDR activation in cultured CMs is sufficient to inhibit CM proliferation, they seem to be dispensable for CM cell-cycle exit and binucleation *in vivo*.

**Fig. 5.**
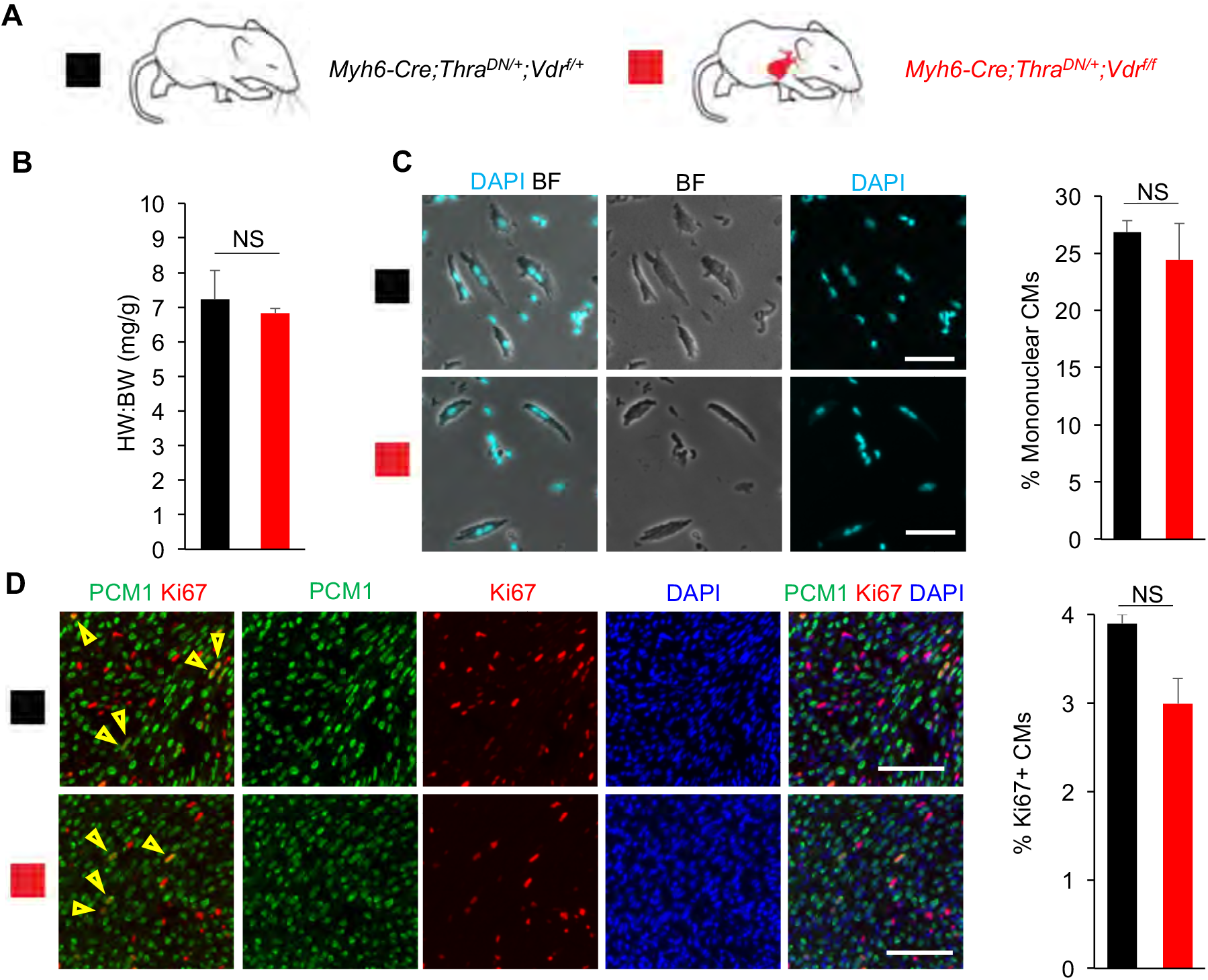
Loss of Vdr is insufficient to alter the *in vivo* proliferative potential retained by thyroid hormone (TH) signaling-deficient CMs. (**A**) Schematic for generating *Myh6-Cre;Thra*^*DN*/+^;*Vdr*^*f/f*^ mice with CM-specific deletion of floxed endogenous VDR and CM-restricted expression of a dominant negative (DN) TH receptor alpha. (**B**) Heart weight (mg) to body weight (g) ratio (HW:BW) did not differ between *Myh6-Cre;Thra*^*DN*/+^;*Vdr*^*f/f*^ mice and *Myh6-Cre;Thra*^*DN*/+^;*Vdr*^*f/f*^ controls at P14. (**C**) Mononuclear CM percentage did not differ between *Myh6-Cre;Thra*^*DN*/+^;*Vdr*^*f/f*^ mice and *Myh6-Cre;Thra*^*DN*/+^;*Vdr*^*f/f*^ controls at P14. (**D**) No difference in the number of Ki67^+^ CMs was observed between *Myh6-Cre;Thra*^*DN*/+^;*Vdr*^*f/f*^ mice and *Myh6-Cre;Thra*^*DN*/+^;*Vdr*^*f/f*^ controls at P14. Values are reported as mean ± SEM. (n=3) NS, not significant. \**p* < 0.05. Scale: (C) 50 μm (D) 100 μm.

## DISCUSSION

Mammalian cardiomyocytes undergo postnatal cell-cycle exit and polyploidization when they lose regenerative potential^7,8,33–35^. The intrinsic and extrinsic molecular drivers of this process have been under intensive studies^4,36–40^. Our observation that both glucocorticoids and vitamin D inhibit proliferation of primary neonatal mouse CMs *in vitro* is consistent with previous reports in culture^21,22,28,29,41^. GR activation inhibits neonatal rat CM proliferation and increases CM binucleation^21,22^. Vitamin D treatment reduces expression of c-myc and proliferating cell nuclear antigen, and blocks cell proliferation in rat CM culture^41^.

Our *in vivo* analysis, however, did not reveal a significant role for either receptor in regulating postnatal CM proliferative potential loss: neither CM nucleation nor CM proliferation was influenced by either GR or VDR loss in P14 mice. As such, while GR and VDR signaling are sufficient to inhibit CM proliferation in culture, we did not observe their regulatory influence on CM proliferative potential to be determinative *in vivo*. Moreover, we didn’t observe cardiac hypertrophy of these mutant mice at P14, suggesting that the hypertrophic phenotypes described in both mutant mice^31,32^ develop later in the postnatal life. Nevertheless, the discrepancy between our *in vitro* and *in vivo* data could be explained if there are *in vivo* factors not present *in vitro* whose regulatory influence over CM proliferative potential is strong enough to mask any contributions from either GR or VDR signaling.

One such factor could be thyroid hormones (THs). We previously established a potent inhibitory role for TH signaling over CM proliferative potential^11^. Still, this still left open the possibility that GR or VDR signaling may alter the observed effects of loss of TH signaling on CM proliferative potential. When we analyzed mice both expressing dominant negative TH receptor alpha and lacking either GR or VDR in their CMs, no significant effects of either GR or VDR loss were detected. In other words, the regulatory influence of TH over CM proliferative potential is strong enough to overpower any contributions from either GR or VDR. Further investigation is needed to explore the molecular and endocrine factors that act in parallel to and downstream of thyroid hormone to establish vertebrate cardiac regenerative potential. In sum, our results suggest that GR and VDR don’t function as major triggers of postnatal CM cell-cycle withdrawal and binucleation *in vivo.* Possible explanations could involve the physiological levels of these hormones that might not be high enough, and possible other regulatory mechanisms that prevent the action of GR and VDR *in vivo*.

It is worth noting that VDR signaling may have distinct regulatory roles across vertebrates. It is reported recently that in zebrafish, VDR activation stimulates CM proliferation, while VDR antagonism reduces CM proliferation and compromises cardiac regeneration^30^. This sharply contrasts with VDR signaling in mammals, which seems to exert broadly anti-proliferative effects ^27–29^, as corroborated by our *in vitro* observations. It would be interesting to unravel the differential roles of VDR and underlying mechanisms on cell cycle activity across phylogeny.

## ACKNOWLEDGEMENTS

We thank Rachel B. Bigley, Alexander Y. Payumo, and Kentaro Hirose for help breeding and genotyping the mutant mice in our experiments.

## Author contributions

S.C. and D.L. performed experiments. S.C. and D.L. collected and analyzed data. S.C., D.L., and G.N.H. contributed to discussions. S.C. and G.N.H. designed experiments and wrote the manuscript.

## MATERIALS & METHODS

### Animals

Mouse protocols were conducted in accordance with the Institutional Animal Care and Use Committee (IACUC) of the University of California, San Francisco. CD-1 (Charles River), *Myh6-Cre* (20), *TRα*^AMI/AMI^ (here named *Thra*^DN/DN^) (12), *Nr3c1*^*f/f*^ (here named *Gr*^*f/f*^) (8), and *Vdr*^*f/f*^ (9) mouse lines were maintained according to the University of California, San Francisco institutional guidelines. All mice were 14 days old when analyzed for *in vivo* experiments: cardiomyocyte nucleation and proliferation. Primary cardiomyocytes for *in vitro* experiments were cultured from P1 CD-1 neonatal mice of mixed sex. For all experiments, both male and female mice were used and no gender difference was observed.

### Reagents

The following reagents were used: rabbit anti-PCM1 (1:2000) (Sigma HPA023370), rabbit anti-PCM1 (H262) (1:200) (Santa Cruz Biotechnology SC-67204), Alexa Fluor 488 donkey anti-rabbit IgG (Molecular Probes 1874771) (1:500), Rat anti Ki-67 monoclonal antibody (SolA15), Southern Biotech Dapi-Fluoromount-G Clear Mounting Media (Southern Biotech 0100-20), 5-Ethynyl-2-deoxyuridine (Santa Cruz Biotechnology SC-284628), Click-it EdU imaging kit (ThermoFisher Scientific C10337), Collagenase Type II (Worthington LS004177), Ethyl carbamate (Alfa Aesar AAA44804-18), calcitriol (Sigma-Aldrich 32222-06-3), alfacalcidol (Sigma-Aldrich L91201381), dexamethasone (TCI America TCI-D1961-1G), hydrocortisone (Alfa Aesar AAA16292-03), corticosterone (Enzo 89149-746), sodium chloride (Sigma-Aldrich S9888-10KG), potassium chloride (Sigma-Aldrich P9541-1KG), monopotassium phosphate (Fisher 5028048), sodium phosphate dibasic heptahydrate (Fisher S25837), magnesium sulfate heptahydrate (FisherS25414), HEPES (Sigma-Aldrich H4034-100g), sodium bicarbonate (EMD EM-SX0320-1), taurine (Sigma-Aldrich T8691-100G), biacetyl monoxime (Sigma-Aldrich B0753-100G), glucose (Amresco 97061-164), EGTA (Amresco 0732-288), protease XIV (P5147-100MG), fetal bovine serum (JRS CCFAP004), calcium chloride (Amresco 97061-904), primocin (Invivogen NC9141851), DMEM (CCF CCFAA005).

### Methods

#### Generating mouse lines

Mice heterozygous for the *Tg(Myh6-Cre)2182Mds* transgene (here referred to as “*Myh6-Cre*”) were bred with mice homozygous for the *Nr3c1*^*f/f*^ allele (here named *Gr*^*f/f*^) to generate *Myh6-Cre;Gr*^*f*/+^ mice. These *Myh6-Cre;Gr*^*f*/+^ mice were then bred with *Gr*^*f/f*^ mice to generate mutant *Myh6-Cre;Gr*^*f/f*^ mice and littermate controls of the following genotypes: *Gr*^*f*/+^, *Gr*^*f/f*^, and *Myh6-cre;Gr*^*f*/+^. Similarly, mice heterozygous for *Myh6-Cre* were bred with mice homozygous for the *Vdr*^*f/f*^ allele to generate *Myh6-Cre;Vdr*^*f*/+^ mice. These *Myh6-Cre;Vdr*^*f*/+^ mice were then bred with *Vdr*^*f/f*^ mice to generate mutant *Myh6-Cre;Vdr*^*f/f*^ mice and littermate controls of the following genotypes: *Vdr*^*f*/+^, *Vdr*^*f/f*^, and *Myh6-Cre;Vdr*^*f*/+^.

In parallel, mice heterozygous for *Myh6-Cre* were bred with mice homozygous for the *Thra*^*DN*^ allele to generate *Myh6-Cre;Thra*^*DN*/+^ mice that were subsequently bred with either mice homozygous for the *Gr*^*f/f*^ allele or mice homozygous for the *Vdr*^*f/f*^ allele to generate both *Myh6-Cre;Thra*^*DN*/+;^*Gr*^*f*/+^ mice or *Myh6-Cre;Thra*^DN/+;^*Vdr*^*f*/+^ mice, respectively. The *Myh6-Cre;Thra*^*DN*/+;^*Gr*^*f*/+^ mice were bred with *Gr*^*f/f*^ mice to generate experimental *Myh6-Cre;Thra*^*DN*/+;^*Gr*^*f/f*^ mice and control *Myh6-Cre;Thra*^*DN*/+;^*Gr*^*f*/+^ mice. The *Myh6-Cre;Thra*^*DN*/+;^*Vdr*^*f*/+^ mice were bred with *Vdr*^*f/f*^ mice to generate experimental *Myh6-Cre;Thra*^*DN*/+;^*Vdr*^*f/f*^ mice and control *Myh6-Cre;Thra*^*DN*/+;^*Vdr*^*f*/+^ mice.

#### Analysis of cardiomyocyte nucleation

Ventricular tissues were fixed in 3.7% paraformaldehyde for 48 hours followed by incubation in 50% w/v potassium hydroxide solution overnight. After a brief wash with PBS, tissues were gently crushed to release dissociated cardiomyocytes. Cells were further washed with PBS three times, deposited on slides, and then allowed to dry out completely. Nuclei were stained with 4’,6-diamidino-2-phenylindole (DAPI). To determine cardiomyocyte nucleation, images of spotted cardiomyocytes were analyzed in Photoshop. The number of mononuclear, binuclear, and polynuclear cardiomyocytes was determined manually using the Count Tool. At least 130 cardiomyocytes per each sample were analyzed.

#### Neonatal cardiomyocyte isolation and culture

Freshly dissected ventricles from P1 CD-1 strain neonatal mice were cut into 4 pieces, and then washed with Perfusion Buffer (12 mM sodium chloride, 1.5 mM potassium chloride, 60 μM monopotassium phosphate, 60 μM sodium phosphate dibasic heptahydrate, 120 μM magnesium sulfate heptahydrate, 1mM HEPES, 4.6 mM sodium bicarbonate, 30 mM taurine, 10 mM biacetyl monoxime, 5.5 mM glucose, and 400 μM EGTA). Cardiomyocytes were isolated in Digestion Buffer (2 mg/mL collagenase II and 500 μg/mL protease XIV in perfusion buffer) for 2h at 37℃ during agitation. Isolated cardiomyocytes were cultured in Plating Media (5% FBS and 100 μg/mL primocin in DMEM) and plated on a 96-well cell culture plate at a density of ~25 thousand cells/well.

#### Neonatal cardiomyocyte *in vitro* immunohistochemistry

24 hours after plating, old Plating Media was changed and 5 μM EdU Plating Media containing each drug of interest at the requisite concentration was added to each treatment well. (Control wells received Plating Media with 5 μM EdU but no dissolved drugs.) 48 hours after drug addition, Plating Media was removed and each well was immediately washed with 100 μL PBS, fixed with 3.7% PFA for 15 minutes at room temperature, permeabilized in 0.2% Triton X-100 in PBS (PBST), blocked in 5% normal donkey serum (NDS) in PBST for 1 hour at room temperature, and incubated with primary antibodies in PBST overnight at 4°C. After primary antibody incubation, sections were incubated in their corresponding secondary antibody for 2 hours at room temperature, and the Click-it EdU imaging kit was used to visualize EdU via conjugation to sulfo-Cyanine 5-azide dye (Lumiprobe A3330). In all samples, nuclei were visualized by staining with DAPI.

#### P14 cardiomyocyte *in vivo* immunohistochemistry

At P14, experimental mice were anesthetized by injection of 20% ethyl carbamate in 1X PBS, their hearts were freshly excised, soaked briefly in 30% sucrose, and then embedded in O.C.T. Compound (Tissue Tek, cat#4583) and flash frozen on a metal block cooled by liquid nitrogen. Embedded samples were then sectioned with a Leica CM3050S to 5 μm thickness. Tissue sections were then fixed in 3.7% paraformaldehyde for 15 min at room temperature, permeabilized in 0.2% PBST, blocked in 5% NDS in PBST for 1 hour at room temperature, and incubated with primary antibodies in PBST overnight at 4°C. After primary antibody incubation, sections were incubated in their corresponding secondary antibody for 2 hours at room temperature. And mounted in DAPI to visualize nuclei.

#### Statistical analysis

The number of samples per each experimental condition is listed in the description of the corresponding figure legend. Statistical significance was determined using the one-way ANOVA test (Fig. 1), and Student’s T-test for the rest of the figures.

